# Immunizing small cell lung cancer mice with isoaspartylated Elavl4 after chemotherapy mimics improved survival of anti-ELAVL4 antibody-positive small cell lung cancer patients

**DOI:** 10.64898/2026.08.31.748151

**Authors:** Diego A. Velarde, Joseph A. Valdes, Hannah Lee, Chunli Yan, Sarah Elmalh, Angie D. Moreno, Daniel J. Mullen, Matthew A. Gladstone, Sanika Gulavani, Qi Nie, Ming Li, W. Martin Kast, Ite A. Offringa

## Abstract

**Introduction:** Small cell lung cancer (SCLC) patients have an ∼8% 5-year survival; new therapies are urgently needed. Approximately 15% of SCLC patients have naturally-occurring low-titer antibodies against neuronal ELAVL proteins, associated with improved response to therapy and significantly improved survival. We previously determined that the anti-ELAVL4 response is triggered by isoaspartylation in the unstructured ELAVL4 N-terminal region.

**Methods:** We used a *Tp53*^fl/fl^;*Rb1*^fl/fl^ inducible SCLC mouse model to test whether 1) immunization with isoaspartylated Elavl4 (isoAsp-Elavl4) prior to SCLC induction improves survival in the absence of any other treatment, and 2) immunization with isoAsp-Elavl4 following completion of 3 rounds of cisplatin+etoposide therapy improves survival. Immunizations contained incomplete Freund’s adjuvant with either a recombinant N-terminal fragment of Elavl4 (amino acids 1-117), incubated under isoaspartyl-inducing conditions, or phosphate-buffered saline (used as the negative control, since Elavl4 spontaneously isoaspartylates). Mice were monitored by blinded assessors until euthanasia was indicated.

**Results:** IsoAsp-Elavl4-immunized animals all became immune responsive, and spontaneous anti-isoAsp-Elavl4 antibodies were observed in 7% of the control animals. Kaplan-Meier analyses revealed that pre-SCLC immunization with isoAsp-ELAVL4 in the absence of other treatments did not affect survival. In contrast, immunization of SCLC mice following chemotherapy significantly improved survival.

**Conclusions:** An anti-isoAsp-ELAVL4 response can be actively induced in mice and significantly increases SCLC survival when given following chemotherapy. This indicates that the anti-isoAsp-ELAVL4 immune response can be leveraged to develop new therapies for SCLC patients.

## Introduction

Small cell lung cancer (SCLC) comprises ∼13% of lung cancer cases with a 5-year survival of ∼8%.^1,2,3^ Most patients are diagnosed with extensive-stage SCLC, in which the disease has spread beyond the lungs. There has been little improvement in survival over the past 30 years. Immune checkpoint inhibitors (ICIs) have been recently added to the standard platin-based and etoposide chemotherapy given to extensive stage disease patients, providing a modest survival benefit.^4,5^ Fine-tuning of ICI therapy is showing further incremental gains in patient survival.^5,6^ Recently, SCLC-targeted therapies against delta-like protein 3 (DLL3) have begun to show promise,^5,7^ underlining the importance of investigating SCLC-specific antigens to develop new therapies.

Here we focus on the antigen ELAVL4, originally known as HuD. Approximately 15% of SCLC patients show spontaneous antibodies against one or more neuronal ELAVL proteins, encoded by the *ELAVL 2, 3*, and *4* genes.^8^ These proteins show considerable homology and are expressed throughout the nervous system.^9^ In the lungs, only pulmonary neuroendocrine cells (the principal cell of origin of SCLC^10,11^) express neuronal ELAVL proteins, with ELAVL4 showing the highest expression^12^ (Supplementary Figure 1). Patients with modest levels of anti-ELAVL antibodies show improved response to chemotherapy and significantly improved survival.^13,14^ Less than 1% of SCLC patients develop high titer anti-ELAVL antibodies and associated paraneoplastic encephalomyelitis/sensory neuronopathy (PEM/SN), a debilitating autoimmune disease that causes widespread neuronal destruction and often death.^8^ In these patients SCLC may completely regress, underlining the power of the anti-ELAVL response to combat SCLC.^8,14^

**Figure 1.**
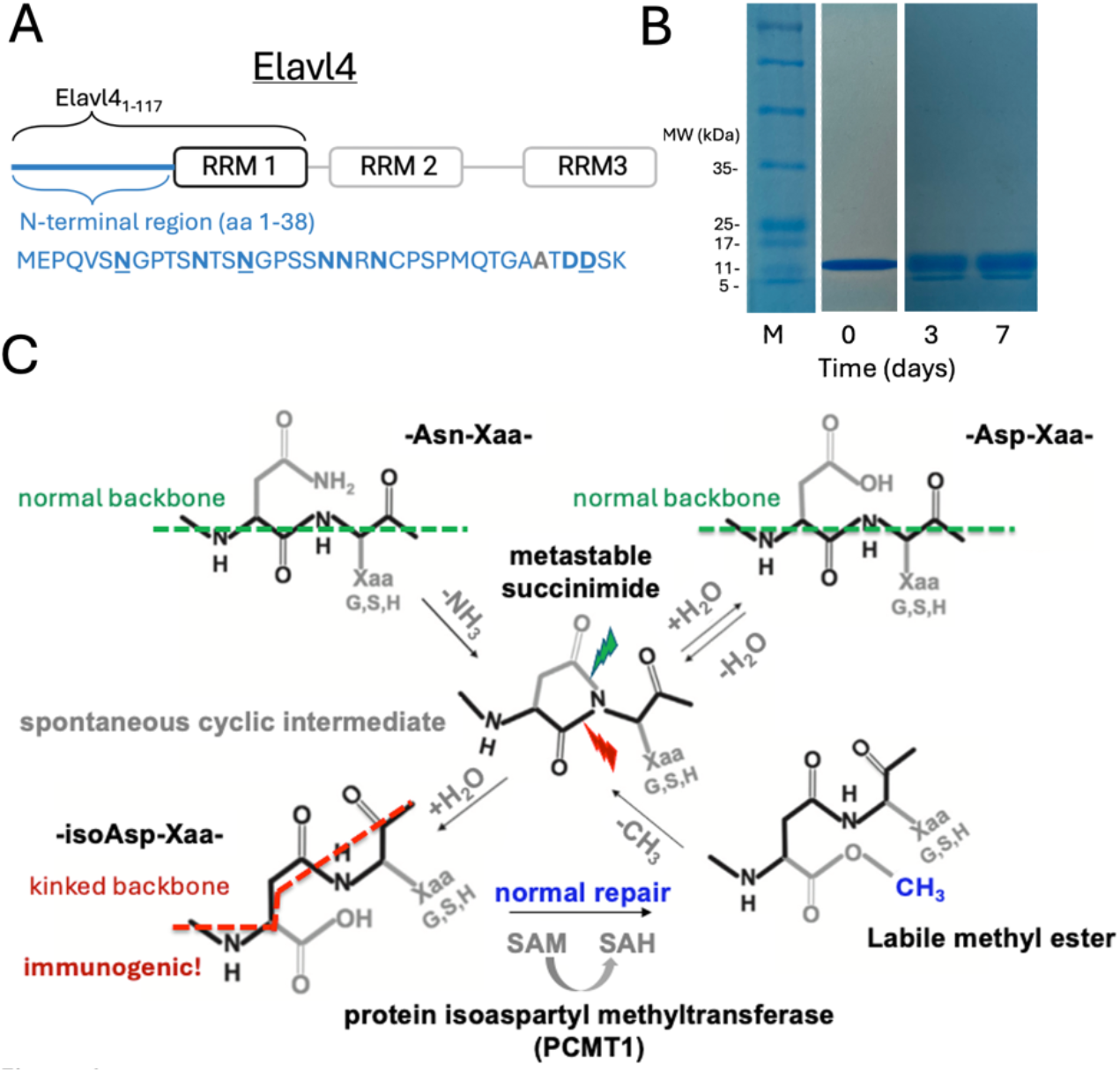
Isoaspartylation in ELAVL4. *(A)* Schematic of Elavl4 RNA Recognition Motif (RRM) domains and N-terminal sequence. The single amino acid difference between mouse and human ELAVL4 (alanine 33) is indicated in bold gray. All Ns and Ds (potential isoaspartylation sites) are in bold blue and canonical isoaspartylation sites (N or D followed by S, G, or H) are underlined. *(B)* Coomassie-stained gel of recombinant Elavl4_1-117_ aged in PBS at 37°C for 0, 3, or 7 days. This results in a characteristic shift upwards of the monomer band and faint higher molecular weight complexes^17,19^. *(C)* Schematic of the mechanism of isoaspartylation and repair.

Serum from a SCLC patient with PEM/SN was originally used to identify ELAVL4 as the antigen^15^ and subsequent work showed that ELAVL proteins are present on the surface of SCLC tumors^16^. We demonstrated that anti-ELAVL4-positive SCLC patient serum samples react specifically with the 38-amino acid unstructured ELAVL4 N-terminal region that lies upstream of the first of its three globular RNA-recognition motif (RRM) domains.^17^ We identified unrepaired isoaspartylation in the N-terminal region (Figure 1) as the trigger of the anti-ELAVL4 response.^17^ Isoaspartylation is a naturally-occurring highly immunogenic post-translational damage that occurs at asparagine and aspartate residues in unstructured regions; it kinks the peptide backbone (Figure 1*C*) and causes protein aggregation.^17,18,19^ In healthy tissue, the enzyme protein-L-isoaspartate O-methyltransferase (PCMT1) repairs isoaspartylation. PCMT1 is ubiquitous but is particularly highly expressed in the nervous system.^20^ *Pcmt1* knockout mice die of seizures soon after birth^20^ and we have shown that in the brains of these mice the N-terminal region of Elavl4 undergoes isoaspartylation.^17^ Recombinant ELAVL4 spontaneously undergoes isoaspartylation *in vitro* at neutral pH and is immunogenic, inducing B- and T-cell responses in mice and stimulating proliferation of human peripheral blood monocytes.^17^ Using an affinity-purified rabbit polyclonal anti-isoaspartylated ELAVL4 serum we previously detected isoaspartylated ELAVL4 in all samples of human SCLC tumors examined.^17^ Together, these observations suggest that isoaspartylated ELAVL4 accumulates in SCLC and constitutes a cancer-specific neoantigen, to which a therapeutic immune response might be raised.

We thus hypothesized that immunization with isoaspartylated-ELAVL4 in the context of SCLC would improve survival. To test this hypothesis, we used a transgenic *Tp53*^fl/fl^;*Rb1*^fl/fl^ inducible SCLC mouse model^21^ that was developed based on the bi-allelic loss of *TP53* and *RB1* seen in ∼ 90% of human SCLC.^22^ The resulting cancer, originating mainly from the pulmonary neuroendocrine (PNE) cells in the lung epithelium^10,11^ recapitulates the histology, metastatic behavior,^21^ and spontaneous anti-ELAVL4 antibodies^23^ of human SCLC. Here we used the *Tp53*^fl/fl^;*Rb1*^fl/fl^ SCLC mouse model to test whether actively inducing an immune response to isoaspartylated Elavl4 through immunization is therapeutic. We show that immunization with recombinant Elavl4_1-117_, incubated under isoaspartyl-inducing conditions (henceforth referred to as isoAsp-Elavl4) induces a prolonged antibody response and, when given following chemotherapy, provides a significant survival benefit. This indicates that the anti-isoAsp-ELAVL4 response may be leveraged to develop new therapies for SCLC patients.

## Materials and Methods

### Recombinant Protein Production

Recombinant Elavl4_1-117_ was produced as described^17^ using endotoxin-negative ClearColi (Lucigen, cat#60810-1). Human and mouse ELAVL4 are fully conserved except for a single threonine (human) to alanine (mouse) change at position 33 (Figure 1*A*). The C-terminally (his)_6_-tagged protein was affinity-purified by adding 400 uL of Ni-NTA agarose (Qiagen, Cat # 30210) to supernatant from 1 L culture that had been pelleted and sonicated in 10 mL TNT buffer (10 mM Tris-HCl pH 8.0, 350 mM NaCl, 0.5% Triton X-100, 10% glycerol, 1 mM Tris(2-carboxyethyl)phosphine), rotated for 1 hour at 4°C, and subjected to 3-minute washes with 5 mL of 20 mM imidazole in TNT and with 3 mL of 50 mM imidazole in TNT, then eluted thrice with 200 uL of 250 mM imidazole in TNT, and twice with 200 uL of 500 mM imidazole in TNT (3 min). The elutions with the highest Elavl4_1-117_ concentration were determined using SDS-PAGE, were combined and buffer-exchanged into phosphate-buffered saline (PBS) pH 7.4 using Amicon Ultra Centrifugal Filters 3K MWCO (Millipore, cat# UFC9010). Protein was quantitated using the Pierce BCA protein assay kit (ThermoFisher, cat # A55865). One batch of protein was used for all immunization experiments. The protein was filter sterilized using a 0.22-micron filter. Half of the protein batch was subjected to isoaspartylation by incubation at 37°C for 7 days. Presence and purity of protein samples was confirmed by SDS-PAGE (Figure 1*B*). Protein was stored in aliquots at -20°C until ready for use (this minimizes isoaspartylation of the native protein).

### Mouse Immunization, Monitoring, Chemotherapy and Serum Collection

Animal studies were performed in accordance with the guidelines of the USC Institutional Animal Care and Use Committee (approved protocol IACUC # 20105). Mice were *Trp53*^fl/fl^; *Rb1*^fl/fl^ in an FVB/N background as previously described^21^ and were enrolled following genotype confirmation (*Trp53*^fl/fl^; *Rb1*^fl/fl^). Both male and female mice were included in all experiments and animals were randomly assigned to experimental cohorts. Sample sizes were determined by power analysis based on survival differences observed in anti-Hu–positive SCLC patients,^14^ yielding approximately 80% power to detect survival effects at α = 0.05. Exclusion criteria included evidence of conditions or technical complications unrelated to experimental intervention. No exclusions were made based on study outcome. Animals that experienced non–tumor-related death were retained in overall survival analyses and were annotated accordingly in study records.

Pre-SCLC immunization experiment: Mice were immunized at 10 weeks of age. IsoAsp-Elavl4 immunization was by subcutaneous injection in the flank with 100 uL containing 150 ug of isoAsp-Elavl4_1-117_ emulsified 1:1 in incomplete Freund’s adjuvant (IFA), prepared using the syringe extrusion method. We immunized control animals with 100 uL vehicle (PBS) in IFA. Mice received an identical booster injection two weeks later. SCLC was induced by intratracheally instilling adenovirus carrying Cre recombinase (Ad5CMVCre, see details below) at 15 weeks. Mice were monitored by blinded assessors using a health rubric (adapted from Paster *et al*.,^24^ Supplementary Figure 2) that scores 1) appearance, 2) natural behavior, 3) provoked behavior, and 4) body condition. Mice were scored every two weeks until they became symptomatic (score of 5 out of 11) at which point they were monitored daily until euthanasia was indicated (score of ≤3 out of 11).

**Figure 2.**
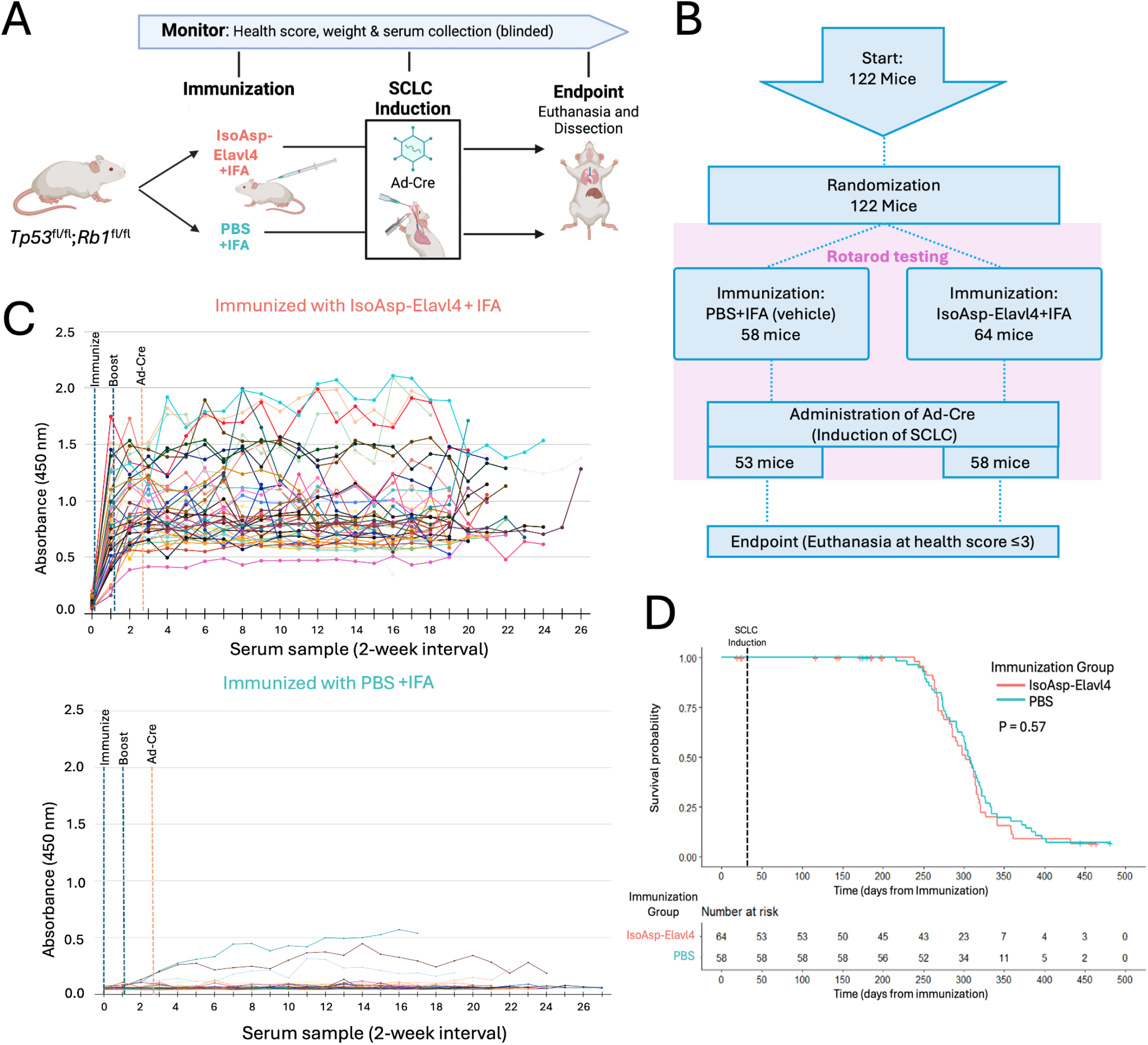
Effect of early isoAsp-Elavl4 immunization in the absence of any other treatment on the survival of mice with SCLC. *(A)* Schematic of experiment using *Trp53*^fl/fl^;*Rb1*^fl/fl^ mice to evaluate the effect of immunization alone on overall survival. *(B)* Flow chart describing the trajectory of animals included in the analysis. IFA=incomplete Freund’s adjuvant. 28 mice were censored for causes of death other than detectable SCLC. In the isoAsp-Elavl4-immunized group, mice were censored due to apparent epileptic seizures 5-12 days after immunization (n=11), apparent seizures after Ad-Cre induction but before SCLC symptoms would be expected (n=5), ascites with no detectable cancer (n=2), a thyroid tumor (n=1), or no detectable tumor at the end of the study (n=3). In the control-immunization group mice were censored due to a thyroid tumor (n=1), a genital infection (n=1) or no detectable tumor at the end of the study (n=4). *(C)* Plot of anti-isoAspElavl4 antibody levels detected in mouse serum. *(D)* Kaplan-Meier survival plot describing the results of the pre-SCLC immunization experiment. Survival is presented from the point of randomization (immunization) until endpoint. Tick marks indicate mice that were censored due death unrelated to SCLC (n=21) or mice that were truncated because they were still alive at the end of the study with no evidence of cancer (n=7). No significant difference was observed between groups relative to SCLC survival (p = 0.57).

Combination cisplatin+etoposide (CE) with immunization experiment: Mice were given Ad5CMVCre at 11 weeks. Symptomatic mice with a health score of 5 were given CE chemotherapy intraperitoneally (day 1: cisplatin (5 mg/kg) and etoposide (10 mg/kg) diluted in PBS; days 2 and 3: etoposide (10 mg/kg) diluted in PBS). Mice received a total of three CE treatments with a 3-week interval between the first day of each treatment cycle. Three weeks after the first day of the last CE cycle mice were immunized with isoAsp-Elavl4 or PBS emulsified in IFA as described above. Two weeks later an identical booster was given.

Longitudinal serum collection began just prior to immunization in the first experiment and following SCLC induction in the CE+immunization experiment and was continued every two weeks for the duration of the mouse’s life. Blood (up to 200 uL), collected from the saphenous vein using Sarstedt 200 uL microvette tubes, was allowed to coagulate at room temperature for 30 minutes and held at 4°C up to 24 hours before centrifugation (3,000 RCF, 5 min). Serum was isolated and stored at -80^°^C until used for analysis.

### RotaRod tests

Mice in the pre-SCLC immunization experiment were tested for adverse motor function and coordination effects before and after immunization using a RotaRod instrument^25^ (Ugo Basile, Cat # 47650). Mice were conditioned on the instrument one week prior to the first data collection. During each run, the initial speed was set to 4 rotations per minute (RPM) with a ramp-up of 1 RPM each 3 seconds until 40 RPM was reached as the final rotation speed. Time (in seconds) to falling was recorded automatically by touch-sensitive pads on the instrument. Mice achieving 500 seconds of running time were stopped manually to avoid overly tiring or stressing the mice. Mice were given 3 trials with 5 min of rest between runs. The device was cleaned with 70% ethanol between individual mice.

### SCLC induction

Mice were administered cyclosporine (20 mg/kg/day) via their drinking water for 1 week prior to and 2 weeks post infection with adenovirus carrying Cre recombinase (Ad5CMVCre, University of Iowa Viral Vector Core). To intratracheally instill Ad5CMVCre, mice were anesthetized with an intraperitoneal injection of ketamine (80 mg/kg) and xylazine (5 mg/kg) and suspended in a supine position to provide access to the trachea. Twenty uL (2.5x10^7^ particle-forming units) of Ad5CMVCre were delivered intratracheally into the lungs using a 2.8-cm polyurethane catheter (0.012”/0.3 mm ID x 0.025”/0.6 mm OD, Access technologies, cat# BC-2P) slipped onto a 29-gauge 0.3 mL insulin syringe. The syringe was filled with 100 uL air, 20 uL virus solution and 50 uL air prior to intubation and the syringe contents were gently delivered into the lungs.

### Detection of antibodies in mouse serum

Anti-isoAsp-Elavl4 antibodies were measured by enzyme-linked immunosorbent assay (ELISA). Recombinant isoAsp-Elavl4_1-117_ (50 uL, 2 ug/mL per well) was incubated overnight at 4°C in a 96-well Immulon 4HBX ELISA plate (VWR cat#62402-972). After three 5-min washes with PBST (PBS, 0.1% TWEEN20) plates were blocked with 200 uL of 3% BSA (Millipore, cat # 81-065-1) in PBS for two hours at room temperature. After three 5-min washes with PBST, 100 uL mouse serum (1:1000 in 3% BSA in PBS) or mouse His-Tag antibody (Proteintech, Cat# 66005-1-Ig, RRID: AB_11232599, diluted 1:1000 in 3% BSA in PBS) were loaded onto the plate and incubated overnight at 4°C. After three 5-min washes with PBST, 100 uL goat-anti-mouse IgG (H+L) HRP secondary antibody (Invitrogen catalogue# 31430, RRID: AB_228307) diluted 1:50,000 in 3% BSA in PBS was added and incubated for two hours at room temperature. After three 5-min washes with PBST plates were developed by adding 50 uL tetramethylbenzidine solution (Themo Scientific Cat# 34029) for 10 min at room temperature, then quenching by the addition of 50 uL 2M sulfuric acid. Plates were read using a ClarioStar^Plus^ plate reader (BMG LABTECH) to quantify absorbance at 450 nm.

### Tumor collection and tissue analysis

Mice with a health score of ≤3 were euthanized by intraperitoneal injection of Euthasol (0.5 g pentobarbital/kg, Covertrus Cat# 11695-4860-1) and subsequent cervical dislocation. A terminal bleed was then withdrawn by cardiac puncture. Lung tumors and metastases were identified by gross *in situ* examination of individual lung lobes, liver, and brain.

### Statistical Analysis

Kaplan-Meier survival analyses were performed in R v4.4.2 using the packages survival v3.8.3 and survminer v0.5.0. Log-Rank test p-values comparing survival distributions between the displayed treatment groups are shown.

### Data Availability Statement

The data generated in this study are available upon request from the corresponding author. Experimental procedures were conducted using established methodologies or standardized laboratory protocols as described. Detailed experimental protocols are available from the corresponding author upon request. Expression profile data analyzed in this study were obtained from the IPF Cell Atlas^12^ (https://www.ipfcellatlas.com) and Gene Expression Omnibus (GEO) (accession: GSE60052).^26^

## Results

### Pre-Immunization with IsoAsp-Elavl4 Does not Improve Survival of SCLC Mice in the Absence of Chemotherapy

We first investigated the effect of inducing an anti-isoAsp-Elavl4 immune response prior to SCLC induction on the survival of the mice in the absence of any other therapy. 122 ten-week-old FVB/N *Tp53*^fl/fl^;*Rb1*^fl/fl^ mice were randomized by individual cages (aiming for approximately equal numbers of males and females) to receive either isoAsp-Elavl4 or vehicle immunization prior to Ad5CMVCre-based SCLC induction (Figure 2*A* and *B*). Because high titer antibodies against ELAVL4 are associated with PEM/SN in humans, we used a RotaRod instrument^25^ to test motor coordination and balance of the isoAsp-Elavl4-immunized and control mice. Analysis of the RotaRod data showed substantial variability between individual mice but did not indicate a significant difference in performance between the isoAsp-Elavl4-immunized and control groups, nor did we observe a sex-based difference (Supplementary Figure 3), suggesting no gross adverse neuro-motor defects due to the immunization. Mice appeared to improve slightly in their ability to stay on the rod over the first six weeks and appeared to show a gradual decline over the next ten weeks, possibly associated with test-associated stress or SCLC development. To avoid unduly stressing the mice as they were developing SCLC, we concluded the RotaRod testing after 16 weeks.

**Figure 3.**
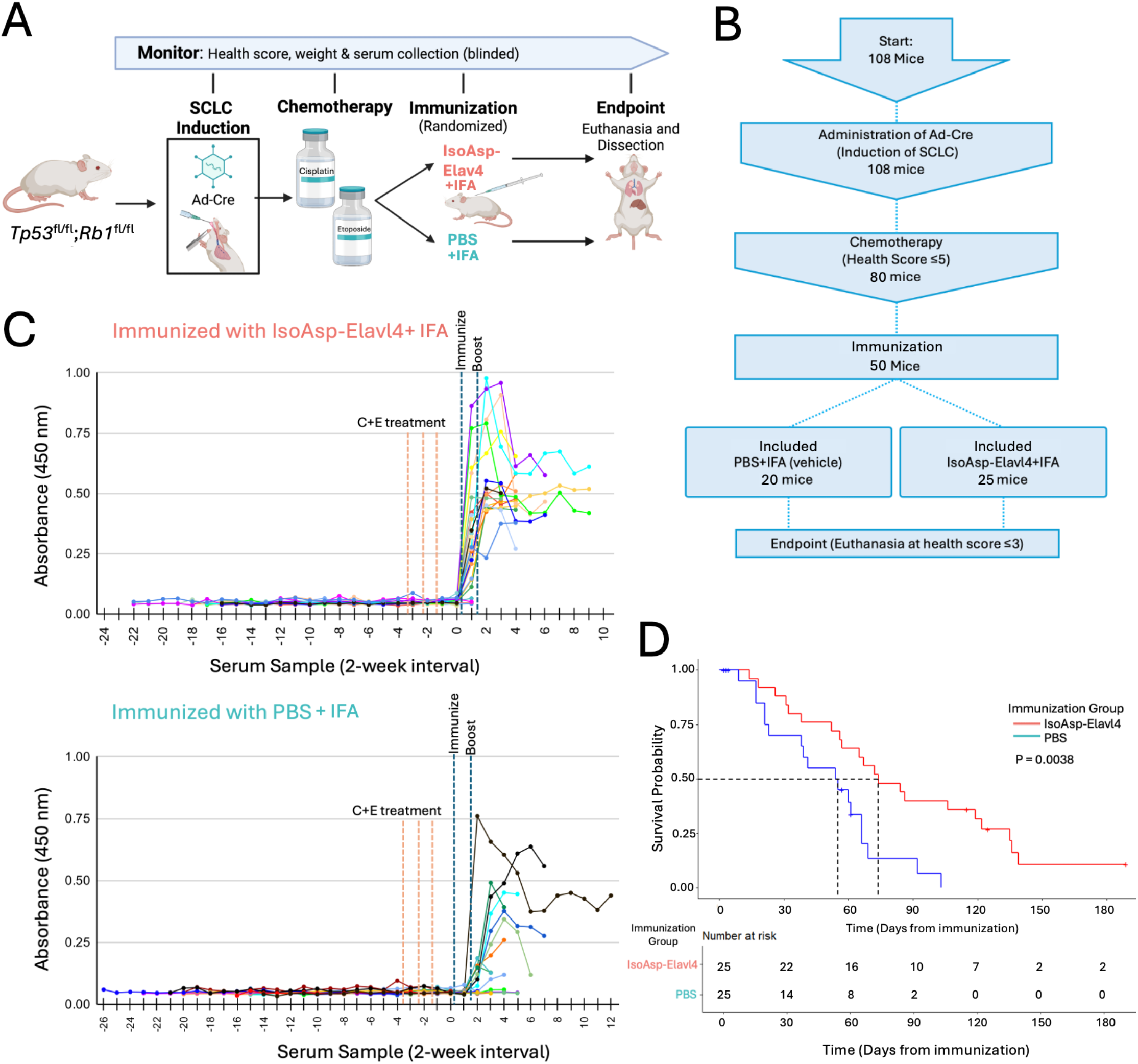
Effect of immunization with isoAsp-ELAVL4 following cisplatin+etoposide (CE) therapy. *(A)* Schematic of experiments and results using *Trp53*^*fl/fl*^*;Rb1*^*fl/fl*^ mice to evaluate the effect of immunization after chemotherapy on overall survival. *(B)* Flow Chart of mice. IFA=incomplete Freund’s adjuvant. 108 Mice received intratracheal instillation of Ad-Cre to induce SCLC. 80 mice survived long enough to receive chemotherapy. 54 mice were randomized to isoAsp-Elavl4 or control-immunization groups. 4 mice died before receiving an immunization (3 isoAsp-Elavl4- and 1 control-immunized) and so 50 mice received an immunization. 5 control-immunized mice died within a week of the boost immunization and were censored (25 isoAsp-Elavl4- and 20 control-immunized mice were included). 4 isoAsp-ELAVL4- and 2 control-immunized mice were truncated and censored from final analysis. *(C)* Plot of anti-isoAsp-Elavl4 antibody levels detected in mouse serum. Serum samples were collected every two weeks *(D)* Kaplan-Meier survival plot describing the results of the CE+immunization experiment. Tick marks represent mice that died before immunization was complete or were truncated at the end of the study and were censored. Survival is presented from the point of randomization (immunization) until endpoint (P = 0.0038).

Longitudinal analyses of mouse serum using ELISA indicated that isoAsp-Elavl4-immunized mice showed a sustained anti-isoAsp-Elavl4 response that varied in magnitude between animals but was consistent over time (Figure 2*C*). Most control-immunized animals showed no reactivity although 4 mice exhibited gradually rising antibody levels possibly related to a naturally occurring response during SCLC development. We previously observed spontaneous anti-ELAVL4 antibodies in a subset of mice with SCLC (14%),^23^ matching what is seen in human patients.

We noted that some female mice had seizures, a phenomenon previously described for FVB/N mice and most commonly occurring in females.^27,28^ In total, 11 female mice from 4 cages died from unknown causes before SCLC could be induced (5-12 days after the boost injection). After Adeno-Cre induction another 5 mice (4 female, 1 male) died before the expected onset of SCLC symptoms. Upon unblinding, we determined that all 16 mice that died were from the isoAsp-Elavl4 immunization group, suggesting that immunization with isoAsp-Elavl4 might have adverse effects. However, independent immunizations aimed at raising antisera and consisting of a total of 5 immunizations, each 2 weeks apart, showed no negative effects or deaths (unpublished results). Indeed, we did not observe such unexplained deaths in an environment where no RotaRod testing was done, thus, a combination of factors may be at play.

Assessors blinded to immunization status monitored mice using a comprehensive health rubric (Supplementary Figure 2*A*). Signs of SCLC appeared 6-8 months after Adeno-Cre induction, as described previously^21^. Mice were euthanized when they reached a health score ≤3 out of the healthy score of 11. Health scores did not show a significant difference between isoAsp-Elavl4 and control-immunized mice (Supplementary Figure 2*B*). Kaplan-Meier analysis indicated no significant difference in survival of mice immunized with isoAsp-Elavl4 compared to control (Figure 2*D*). Within the isoAsp-Elavl4-immunized group, the antibody titer was not related to survival (Supplementary Figure 4*A*). Thus, in the absence of any other therapy, immunization with isoAsp-Elavl4 is insufficient to improve survival of mice that later develop SCLC.

### Immunization with IsoAsp-Elavl4 after Chemotherapy Improves Survival of with Mice SCLC

We next tested whether immunization with isoAsp-Elavl4 following chemotherapy would extend survival of mice with SCLC. We reasoned that the period following chemotherapy would provide a clinically viable opportunity for isoAsp-Elavl4 immunization. We thus instilled adeno-Cre in 108 mice and used the health score to monitor mouse health (Figure 3*A* and *B*). Blood was drawn at 16 weeks of age and every 2 weeks thereafter throughout the animals’ lifetimes to monitor any anti-Elavl4 response (Figure 3*C*). At the onset of SCLC symptoms, reflected by a health score of ≤5, mice were given three cycles of cisplatin+etoposide (CE) therapy, mimicking treatment provided to human patients. In human patients, about one third of patients do not respond to cisplatin (or carboplatin) with etoposide therapy. In our hands, 50% of the mice died before, during, or soon after chemotherapy, suggesting that starting treatment before a health score of 5 is reached may be indicated. Comparison of the survival of control-immunized mice from the first experiment with control-immunized/CE-treated mice from the second experiment indicates that CE treatment trended toward improved mouse SCLC survival but did not reach significance (p = 0.062) (Supplementary Figure 4*B*).

Mice that completed CE therapy were allowed to recover for 3 weeks after the initiation of the last round of CE therapy. 54 mice were randomized to immunization followed by an identical boost two weeks later; however, 4 mice died before receiving an immunization (3 isoAsp-Elavl4- and 1 control-immunized) and so 50 mice received an immunization (25 isoAsp-Elavl4 + IFA and 25 PBS + IFA). 5 control-immunized mice died within a week of the boost immunization and did not survive long enough to meet the inclusion criteria for the immunization protocol which consisted of immunization, an identical booster injection, and survival 1 week post booster injection. The remaining mice (25 isoAsp-Elavl4 + IFA and 20 PBS + IFA) were considered fully immunized and were closely monitored by blinded observers until euthanasia was indicated as soon as a health score of ≤3 was noted. 4 mice from the isoAsp-ELAVL4 and 2 from the control group did not reach the defined endpoint by the conclusion of the observation period and thus were included in the analysis but censored. The survival times were evaluated using Kaplan-Meier analysis. ELISA analysis indicated that compared to control-immunized mice, mice in the isoAsp-Elavl4 immunization group exhibited an anti-isoAsp-Elavl4 response after completing the immunization and boost, though titers varied (Figure 3*C*). Some mice in the control-immunized group also showed an anti-isoAsp-Elavl4 response post-chemotherapy, which may be related to SCLC and/or the response to chemotherapy (Figure 3*C*). Kaplan-Meier analysis showed a significant improvement in survival when comparing isoAsp-Elavl4-immunized *vs*. control mice (p = 0.0038) (Figure 3*D*).

## Discussion

Our results suggest that the anti-isoAsp-ELAVL4 response could be leveraged therapeutically. This could be done in multiple ways. While immunization with isoAsp-Elavl4 appears to show therapeutic potential, it could also result in an autoimmune response targeting the healthy nervous system, as evidenced by the rare SCLC patients with high titer anti-ELAVL4 antibodies who develop PEM/SN. Indeed, the immune response to isoaspartylated proteins has been documented to expand to the native proteins through epitope spreading, a process that results in immune reactivity against protein regions beyond the initial triggering epitope.^29^ Using a RotaRod test, we did not observe neuromotor effects of immunization with isoAsp-Elavl4 protein. However, numerous mice in the pre-SCLC isoAsp-Elavl4 immunization group died from epileptic seizures. Seizures are known to occur in the FVB/N background, particularly in female mice.^27,28^ Because we did not note a similar frequency of seizures and unexplained deaths in the immunization+CE experiment and in unrelated experiments aimed at raising antibodies involving 5 immunizations, each with a booster shot 2 weeks later (not shown), it is possible that stress from the RotaRod in combination with immunization exacerbates negative effects. Previous experiments using Elavl4 protein to immunize mice yielded high titer antibodies but did not cause paraneoplastic disease.^30^ However, these attempts used mice without SCLC and SCLC might affect the propensity for PEM/SN, for example by changes in the blood-brain barrier associated with the cancer, its metastasis, and/or therapy. The risk for paraneoplastic disease suggests that the use of isoAsp-ELAVL-specific monoclonal antibodies could provide a safer therapeutic alternative than immunization. Such antibodies are under development.

Our observation that the increased survival of human SCLC patients with spontaneous anti-ELAVL4 antibodies can be mimicked by isoAsp-Elavl4 immunization following chemotherapy provides a model in which the mechanism by which the anti-isoAsp-ELAVL4 response improves survival can now be teased out. Immunizations of mice with a variety of isoaspartylated proteins induces B-cell as well as a T-cell responses.^31,17^ Numerous mechanisms whereby anti-isoAsp-ELAVL4 antibodies could result in anti-tumor activity can be envisioned, such as complement activation and natural killer (NK) cell activation.^32^ Involvement of T-cells cannot be excluded, although MHC class I expression has been noted to be low or absent in neuronal-type SCLC, which represents ∼80% of SCLC^5,33^; expression profiling of SCLC has led to the classification of SCLC into 4 types, based on expression of 1) *ASCL1*, 2) *NEUROD1* (both considered “neuronal” and accounting for ∼80% of all SCLC), 3) *POU3F2*, and 4) “inflammatory” SCLC (the latter two considered non-neuronal).^34^ Experiments to test the contribution of different immune cell types to therapeutic isoAsp-Elavl4 immunization will be important to gain mechanistic insight into the effects of immunization.

It is notable that immune checkpoint inhibition therapy has been only moderately effective in SCLC and appears to affect mainly non-neuronal SCLC.^5^ Single-cell RNA-seq data has shown that ELAVL4 is expressed in pulmonary neuroendocrine cells,^12^ the most common cell of origin of SCLC,^10,11^ explaining why *ELAVL4* is expressed in most SCLCs. In fact, analysis of expression data from 79 SCLC shows that *ELAVL4* is more frequently expressed in SCLC than DLL3 (Supplementary Figure 5), a protein against which targeted therapies are presently under evaluation.^7^ Just as with DLL3, ELAVL4 has been shown to be present in the surface of SCLC cells.^16^ In the case of DLL3, SCLC-specificity is provided by the increased presence of DLL3 on SCLC cells compared to non-cancer cells. In contrast, isoaspartylated ELAVL4 provides a cancer-specific epitope; isoaspartylation is repaired by PCMT1 in healthy cells. Combining ICI and an anti-ELAVL4 therapy may be especially effective, as it may mobilize the immune system to attack different SCLC subtypes through different mechanisms.

Our experiments indicate that preventive immunization with isoAsp-Elavl4 is not effective in the absence of chemotherapy. It will be important to determine if CE therapy can activate the therapeutic effect of early isoAsp-Elavl4 immunization. If so, it would indicate that CE may potentiate the anti-tumor isoAsp-Elavl4 response, perhaps by increasing the presence of isoaspartylated Elavl4 on the tumor cells or by improving access of immune cells to the cancer cells. However, anti-ELAVL4 antibodies can arise without CE treatment, as paraneoplastic syndromes in human patients frequently precede the diagnosis of SCLC.^35^

Despite a well-documented correlation between the anti-ELAVL4 response and improved survival in SCLC patients, to date no therapies have been developed based on this relationship. This is the first report describing the ability to induce a therapeutic anti-isoAsp-Elavl4 response in a SCLC mouse model. Our results provide a foundation for developing novel SCLC therapies based on the anti-isoAsp-ELAVL4 response.

## Acknowledgments

Diego Velarde recieved support as a Grad^+^ trainee of CaRE^2^, the Cancer Research Education and Engagement Health Center, which is supported by National Institutes of Health/National Cancer Institute (NIH/NCI) grants U54CA233396, U54CA233444, and U54233465, and from NIH/NCI R21 CA290319-01S1 and a Hastings Center for Pulmonary Research Postdoctoral Fellowship. Joseph Valdes was supported by the Francisco Bravo Medical Magnet High School Science, Technology, and Research (STAR) and Engineering for Health Academy (EHA) programs. Hannah Lee and Sarah Elmalh received support from the USC Provost’s Undergraduate Research Associates Program. Daniel Mullen received support from the John H. Richardson Endowed Postdoctoral Fellowship Award in Oncology Research at the Norris Comprehensive Cancer Center. Matthew Gladstone received support as a Grad^+^ trainee of CaRE^2^ and was supported by a Keck School of Medicine Dean’s Fellowship. W. Martin Kast received support from NIH/NCI R21 CA301276, the Walter A. Richter Cancer Research Chair, and Robert Brennan. Ite Offringa has received support for the present study from the V Foundation, a Wright Foundation Transformative Cancer Grant, NIH/NCI grant R21 CA301276, CaRE^2^ (U54CA233396, U54CA233444, and U54233465), the Norris Comprehensive Cancer Center Core Grant (NIH/NCI P30CA014089), and generous donations by Conya Pembroke and Larry Auerbach. The funding agencies had no role in study design, data collection, data analysis, data interpretation, or writing of the manuscript. The authors would like to thank Minxiao Yang for instruction in mouse dissection, the Norris Comprehensive Cancer Center Cores for Flow Cytometry and Immune Monitoring, Molecular Genomics, and Data Science, and the members of the Offringa and Marconett laboratories for helpful discussions.

