## Supplemental Figure 1, Supplemental Figure 2, Supplemental Figure 3, Supplemental Figure 4, Supplemental Figure 5 for "Immunizing small cell lung cancer mice with isoaspartylated Elavl4 after chemotherapy mimics improved survival of anti-ELAVL4 antibody-positive small cell lung cancer patients"

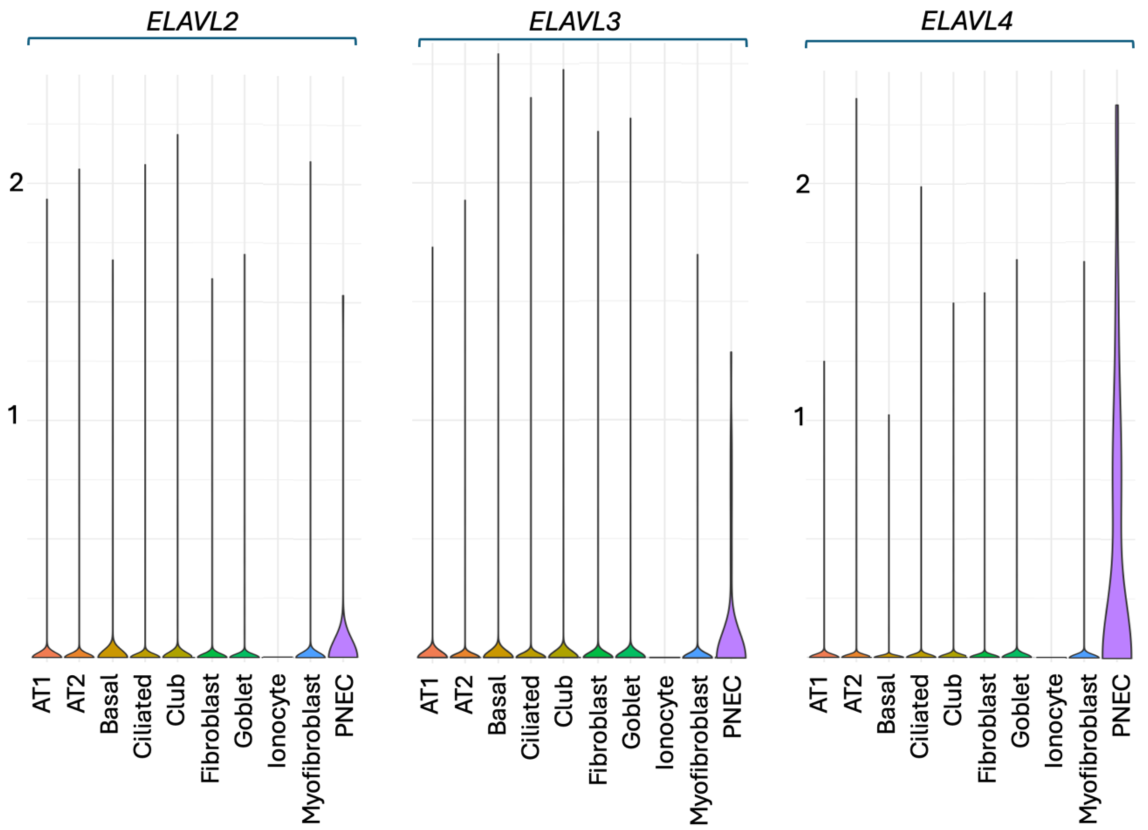


**Supplementary Figure 1.** Single cell RNA-seq data showing expression of neuronal ELAVL genes in human lung cell populations (12) Raw UMI counts were normalized with a scale factor of 10,000 UMIs per cell and subsequently natural log transformed with pseudocount of 1.

A

| **Cage:** | **Mouse #:** | | **Date** |
|---|---|---|---|
| Parameter | Description | Score | |
| Apperance | Normal: bright eyes; shiny; well-groomed hair coat | 2 | |
| | Abnormal: unkept hair coat, dull fur | 1 | |
| | Abnormal: Hunching, piloerection | 0 | |
| Natural Behavior | Normal: Active; interactive in environment | 3 | |
| | Slight decrease in activity; less interactive | 2 | |
| | Abnormal: Pronounced decrease in activity; isolated | 1 | |
| | Abnormal: Possible self mutilation; hyperactive or immobile | 0 | |
| Provoked Behvior | Normal: quickly moves away | 3 | |
| | Slow to move away or exaggerated response | 2 | |
| | Abnormal: moves away after an extended period of time | 1 | |
| | Abnormal: Does not move away or reacts with excessively exaggerated response | 0 | |
| Body Condition Score | Obese | 5 | |
| | Overweight | 4 | |
| | Normal | 3 | |
| | Thin | 2 | |
| | Emaciated | 1 | |
| Overall | Total Points | 0 - 13 | |
| | Weight (g) | | |
| | Comments / Observations | | |

B Pre-SCLC immunization

C CE with immunization

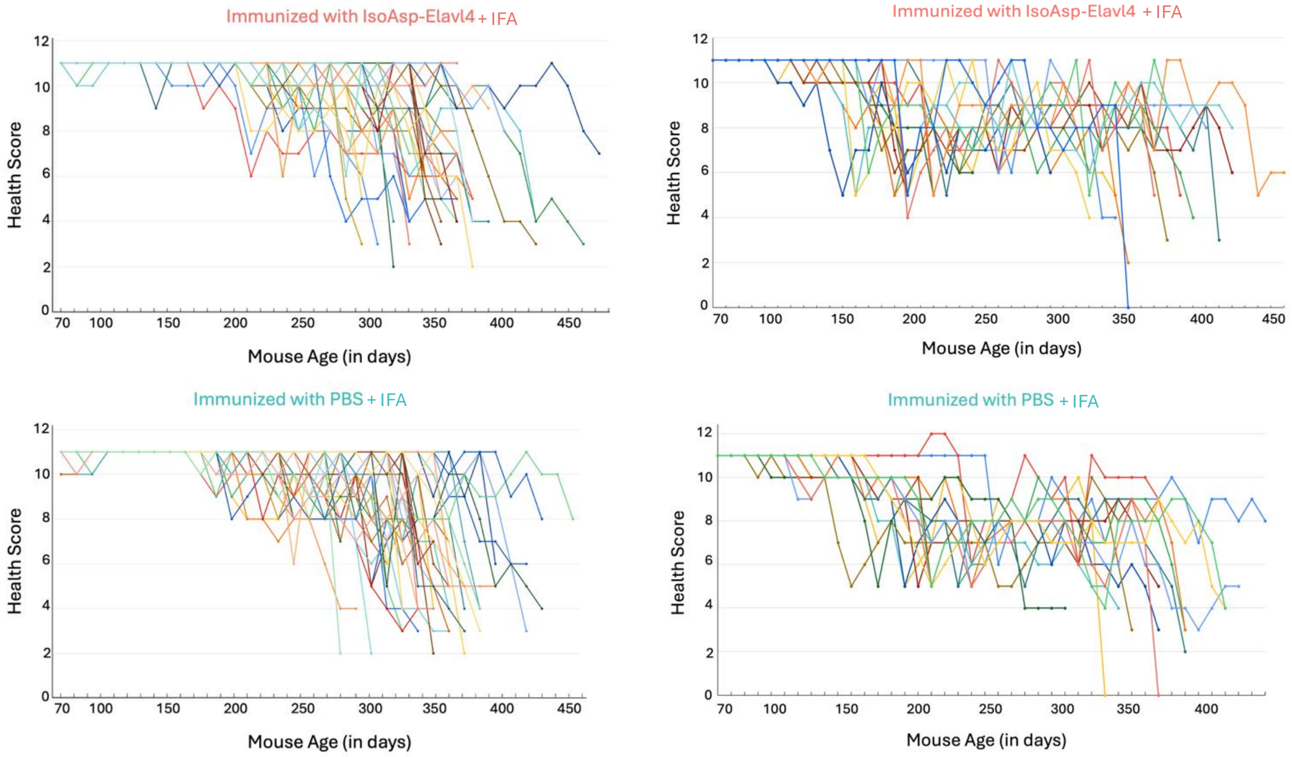


**Supplementary Figure 2.** Health scoring of mice with SCLC to monitor disease progress. *(A)*Scoring rubric (adapted from Paster et al.[23]) with all possible observations indicated. Observations are recorded by an individual blinded to immunization status. A healthy score is 11, mice were treated with CE therapy at a score of ≤5 and euthanized at a score of ≤3. *(B)*Health scores for mice in the pre-SCLC immunization experiment. *(C)*Health scores for mice in the CE with immunization experiment.

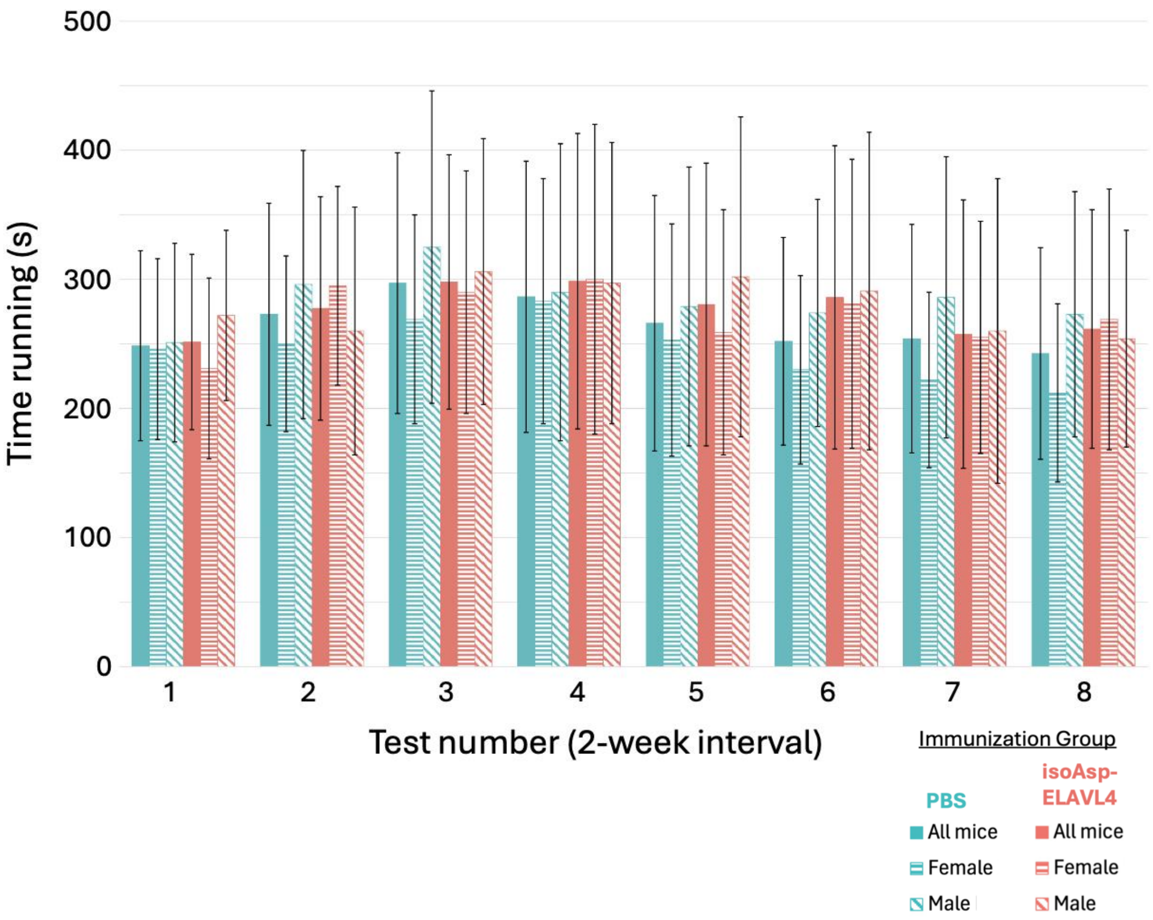


**Supplementary Figure 3.** RotaRod testing results from the pre-SCLC immunization experiment. Mice were tested every two weeks beginning just before immunization and the time to falling was recorded for each mouse. Each time every mouse was tested 3 times separated by 5 min per test.

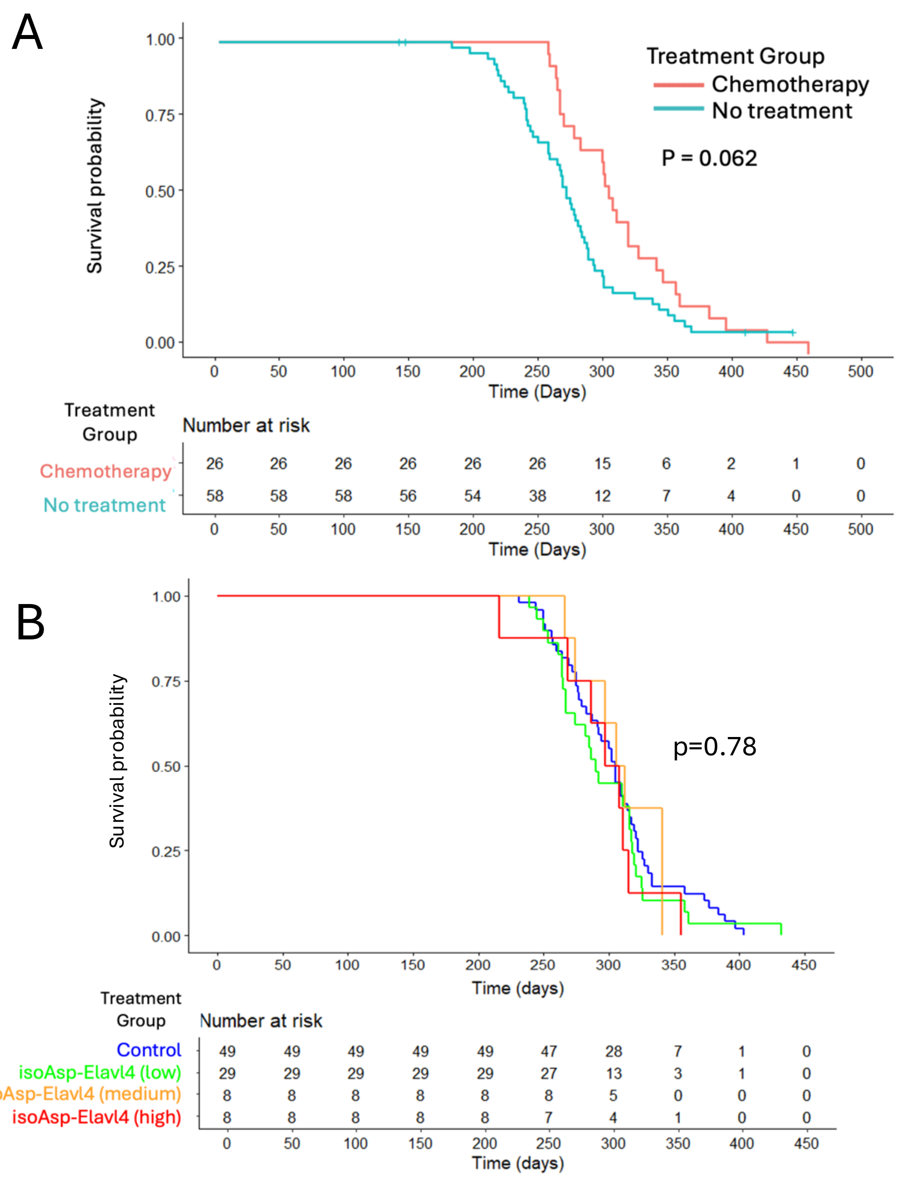


**Supplementary Figure 4.** *(A)*Kaplan Meier survival plot comparing the survival of PBS-immunized mice from the pre-SCLC immunization experiment (blue line) with PBS-immunized/CE-treated mice (red line). Survival is assessed from the point of SCLC-induction until endpoint (euthanasia at health score ≤3). *(B)*Kaplan Meier survival plot comparing the survival of mice from the pre-SCLC-immunization experiment. The isoAspElavl4-immunized mice are stratified based on antibody levels ($ABS_{450}$ value: 1 > low, 1.3 > medium > 1, high > 1.3).

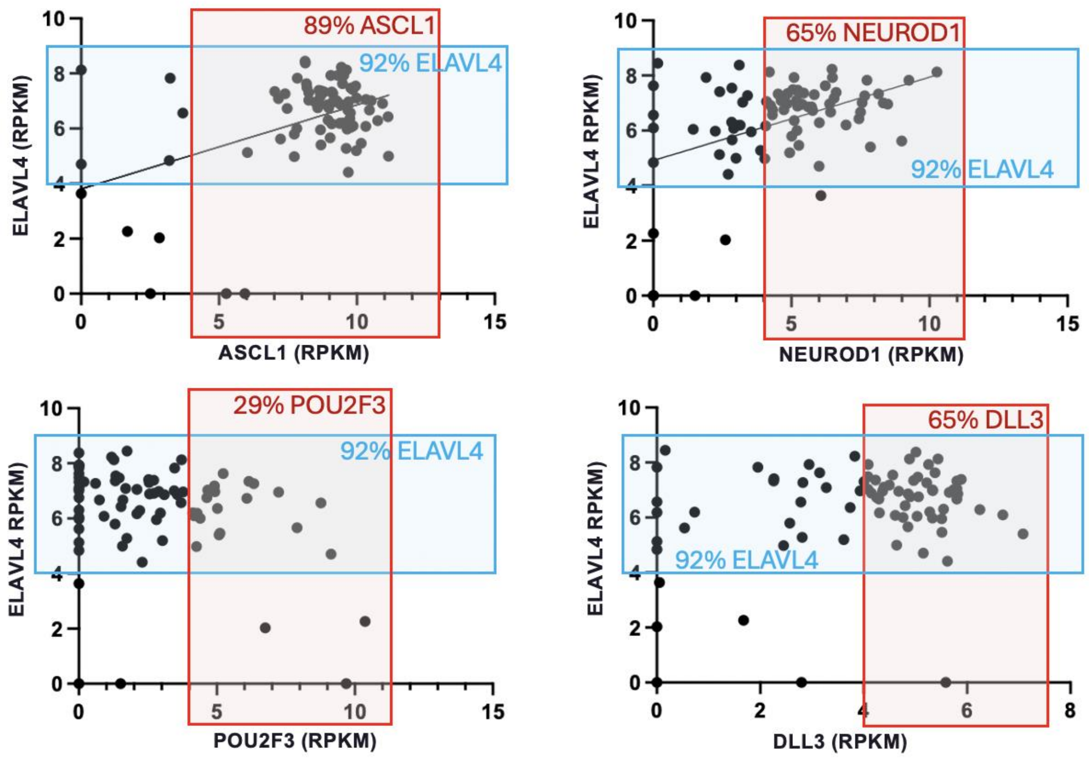


**Supplementary Figure 5.** Scatter plots showing comparative RNA-seq data in reads per kilobase per million mapped reads (RPMK) from 79 SCLC (34) indicating that moderate to high expression (RPKM>4) of ELAVL4 is seen in 92% of SCLC (blue box) capturing most neuronal ($ASCL1^+$ and $NEUROD1^+$) SCLC as well as "non-neuronal" $POU2F3^+$ tumors. In comparison with DLL3, ELAVL4 is expressed in a higher percentage of SCLC (92% vs 68%) than DLL3 (red box bottom right).
